# Generalized dynamics of cooperating bacteria

**DOI:** 10.1101/2025.05.08.652847

**Authors:** Jana C. Massing, Thilo Gross, Justin D. Yeakel, Ashkaan K. Fahimipour

**Affiliations:** MARUM Center for Marine Environmental Sciences, University of Bremen, Bremen, DEU; Department of Marine Glycobiology, Max Planck Institute for Marine Microbiology, Bremen, DEU; Helmholtz Institute for Functional Marine Biodiversity, Oldenburg, DEU; Alfred-Wegener Institute, Helmholtz Centre for Polar and Marine Research, Bremerhaven, DEU; Institute for Chemistry and Biology of the Marine Environment (ICBM), Carl-von-Ossietzky University, Oldenburg, DEU; School of Natural Sciences, University of California, Merced, CA, USA; Santa Fe Institute, Santa Fe, NM, USA; Department of Biological Sciences, Florida Atlantic University, Boca Raton, FL, USA; Center for Complex Systems, Florida Atlantic University, Boca Raton, FL, USA

## Abstract

Bacterial cooperation involves the exchange of metabolites, which can range from costless byproducts of metabolism to intentionally produced and costly molecules. These interactions occur across spatial scales, from direct cell contact to the diffusion of metabolites in the environment. Due in part to this variety of interaction modes, the impacts of mutualism on bacterial community dynamics remain unclear. Using generalized models, we derive conditions for the onset of different dynamical behaviors in communities of bacterial cooperators across these scenarios. These include exchanges of low-cost metabolites, costly cross-feeding, and cases where bacteria produce either most or only a small fraction of available metabolites. Stability depends strongly on metabolic production costs and the balance between metabolite uptake and production. We further show that perturbations to bacteria have larger impacts than to metabolites. Finally, we demonstrate that spatial metabolite diffusion drives pattern formation, emphasizing the links between local stability and spatial structures in bacterial cooperation.

## Introduction

Bacteria release a wide variety of metabolites — products of metabolic reactions — that are often absorbed and used by other bacteria. These metabolite-based interactions shape community composition, ecosystem function, and stability [1, 2]. By outsourcing the production of essential metabolites such as amino acids, lipids, vitamins, cofactors, and signaling molecules [3, 4, 5, 6], bacteria form metabolite trading networks that influence community dynamics [7, 8, 9]. A vast majority of sequenced bacterial genomes lack the ability to independently synthesize all essential molecules, indicating their reliance on outside sources, including other bacteria, for metabolite supplies [10]. This suggests that metabolite sharing underpins widespread commensal and mutualistic interactions in ecosystems and macrobiotic hosts [6, 11, 12]. Understanding how the trade of chemical intermediaries impacts bacterial persistence is therefore crucial for explaining bacterial community dynamics, and for guiding microbiome engineering in human, agricultural, and natural systems [13, 14, 15].

One challenge in developing a general theory of bacterial mutualisms is the diversity of interaction modes. Bacteria can exchange metabolites that vary widely in production cost, ranging from costly, intentionally synthesized compounds [3] to incidental waste byproducts of metabolism [16]. In addition, these interactions occur across spatial scales [6]. At small scales, mutualists may interact through direct contact, such as within biofilms or along intercellular nanotubes [3, 12]. At larger scales, metabolites may leak or diffuse into the surrounding environment, acting as public goods accessible to distant cells [6, 17, 18].

Despite this ecological diversity, a common feature of mutualistic interactions is a tendency to generate positive feedbacks that destabilize communities. Although mutualisms can provide significant benefits to their partners, they also amplify perturbations, leading to runaway growth or mutually reinforced collapses. As a result, mutualism is typically predicted to destabilize ecological communities unless specific conditions are met [19, 20, 7, 8]. In practice, destabilization often occurs rapidly: mutualistic interactions that escalate can drive exponential population growth until limited by external factors such as resource depletion. Thus, mutualisms observed in real systems are invariably counterbalanced by negative feedbacks, often through competition. The resulting interplay of positive and negative forces can produce remarkably stable communities [21].

Understanding the conditions under which mutualistic systems remain stable requires a dynamical systems perspective, grounded in both ecological and physical theory. Models integrating statistical physics, nonlinear dynamics, and complex systems theory have indeed provided key insights into the stability of microbial interactions. Generalized Lotka–Volterra-type models, combined with random matrix theory, suggest that cooperative interactions tend to destabilize microbial communities unless counterbalanced by competition [19]. Extensions of the classic consumer-resource framework [22, 23, 24] reveal that explicitly modeling resource mediation changes the predicted conditions for stability [7, 8]. Incorporating metabolic processes into these models further shows that variation in metabolic rates and resource distributions can profoundly shape microbial community dynamics [25].

However, existing modeling frameworks face important limitations when applied to microbial mutualisms. Generalized Lotka–Volterra models, while valued for their simplicity and flexibility in representing diverse interaction types, cannot account for the metabolic resource-mediated dynamics that are fundamental to microbial interactions [26, 25, 27]. Consumer-resource models address this limitation explicitly but often depend on numerous unknown parameters [28]. Because most bacterial species remain uncultured [29, 30], our understanding of the precise rate laws and functional forms governing microbial interactions is highly limited. Commonly used functions such as Holling [31], Hill [32], and Monod [33] forms capture basic biological constraints like resource uptake and growth [7, 8, 25, 34, 35, 36]. Yet, functions with similar qualitative shapes can differ markedly in their effects on system stability [37, 38]. Moreover, estimating model parameters often requires extensive time series data and fitting procedures, which are rarely feasible even for simple microbial communities [39].

This uncertainty in interaction forms and parameters limits our ability to predict the dynamics of microbial communities across scales [18, 40]. To address these challenges, we apply *generalized modeling* [41, 42], a framework that avoids assuming specific functional forms and allows the exploration of dynamics across diverse interaction modes. Using generalized models, we investigate a common motif in bacterial communities: cross-feeding between two species, where each produces and releases metabolites that support the other’s growth. Given the prevalence and destabilizing potential of this interaction, understanding its dynamics across variation in production costs and spatial scales is crucial for predicting the behavior of larger microbial networks. By focusing on this foundational case, we uncover principles that scale up to influence the stability and functioning of larger microbial ecosystems. These insights offer a foundation for guiding interventions in human, agricultural, and environmental microbiomes.

### Stability of local cooperative metabolite exchange

To analyze the stability of bacterial cross-feeding systems across interaction modes, we applied the generalized modeling framework [41, 42]. Generalized models allow investigation of local dynamics without committing to specific functional forms, making them well-suited for systems with high uncertainty in interaction structure.

Consider a simplified community motif of two bacterial species, *X* and *Y*, each producing and releasing a metabolite (*A* or *B*) that the partner uses for metabolism and growth (Fig. 1). Dynamics of bacterial and metabolite abundances are governed by a four-dimensional system:

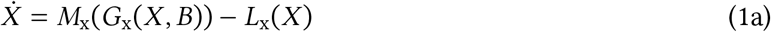

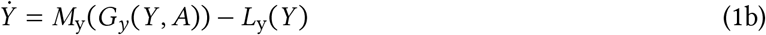

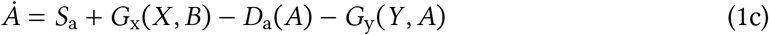

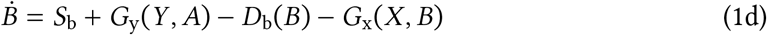

**Figure 1:**
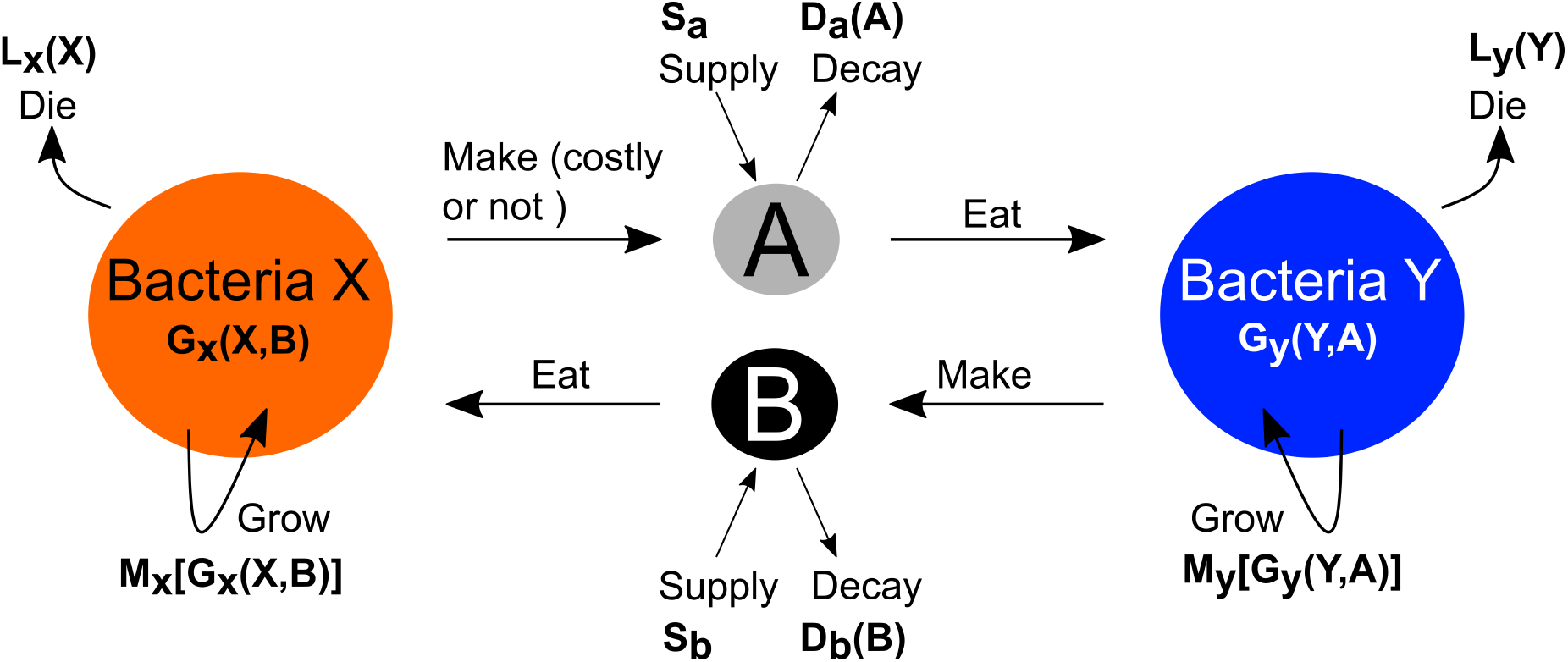
Community motif. Cross-feeding of two bacterial species X, Y on two types of metabolites A,B. Bacterial and metabolite gain and loss processes with corresponding unspecified functions and descriptions of the process they represent.

Here, the function *M* denotes biomass growth, *G* is bacterial metabolic throughput (i.e., uptake, transformation, release), and *L* and *D* denote loss terms for bacteria and metabolites, respectively. The terms *S*_*a*_ and *S*_*b*_ represent exogenous metabolite supply rates.

In conventional modeling, the next step would involve assigning specific functional forms to each process, solving for steady states, and calculating the Jacobian to assess their stability. Generalized modeling diverges from this approach by recognizing that stability can be evaluated without fully specifying the underlying functions; one only needs to parameterize the Jacobian directly, which requires far less information [42]. Instead of defining explicit forms for *M, G, L*, and *D*, we normalize all variables and process rates around an unknown steady state [*X*^⋆^, *Y* ^⋆^, *A*^⋆^, *B*^⋆^], and express the system in terms of normalized rates and elasticities (*i.e*., logarithmic derivatives evaluated at steady state). This yields a fully parameterized Jacobian that captures the stability of any system conforming to the structure of Eq. 1 (see *Methods*).

Local stability is governed by the Jacobian matrix **P**, which describes the linearized dynamics near a steady state. For variables *n* ∈ {*X, Y, A, B*}, its elements are defined as 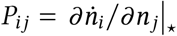, where the ⋆ indicates evaluation at steady state. In the generalized modeling framework, this Jacobian is parameterized using three biologically interpretable sets of quantities: *scale* parameters, which describe turnover rates; *branching* parameters, which capture the relative contribution of gain and loss processes; and *elasticities*, which quantify local nonlinearities via logarithmic derivatives [42, 43].

Scale parameters *α*_*n*_ represent the turnover rate of each variable, or the rate at which biomass or metabolite pools are replenished and lost at steady state. For example,

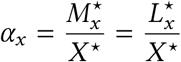

is the *per capita* turnover rate of bacteria *X*, where 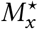 and 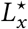 are the steady-state growth and death rates. Analogous parameters *α*_*y*_, *α*_*a*_, and *α*_*b*_ describe turnover for species *Y* and metabolites *A* and *B*.

Branching parameters describe how these gains and losses are distributed across different processes. For instance, the fraction of metabolite *A* lost through decay is

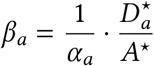

and the fraction gained from external supply is

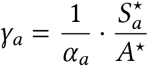

The complementary fractions representing bacterial uptake or production are given by 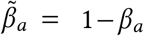 and 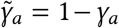. These parameters allow the model to capture a broad range of biologically plausible regimes, from entirely self-supplied metabolites to fully exogenous inputs.

Elasticity parameters quantify local sensitivity of each unspecified function through logarithmic derivatives [44]. For example, the elasticity of *X*’s mortality rate to its own density is

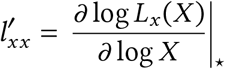

which reflects the nonlinearity of the mortality function near the steady state. Elasticities are dimensionless and defined for all relevant processes in the system and characterize the local curvature or saturation behavior of biological functions (Table 1).

**Table 1:**
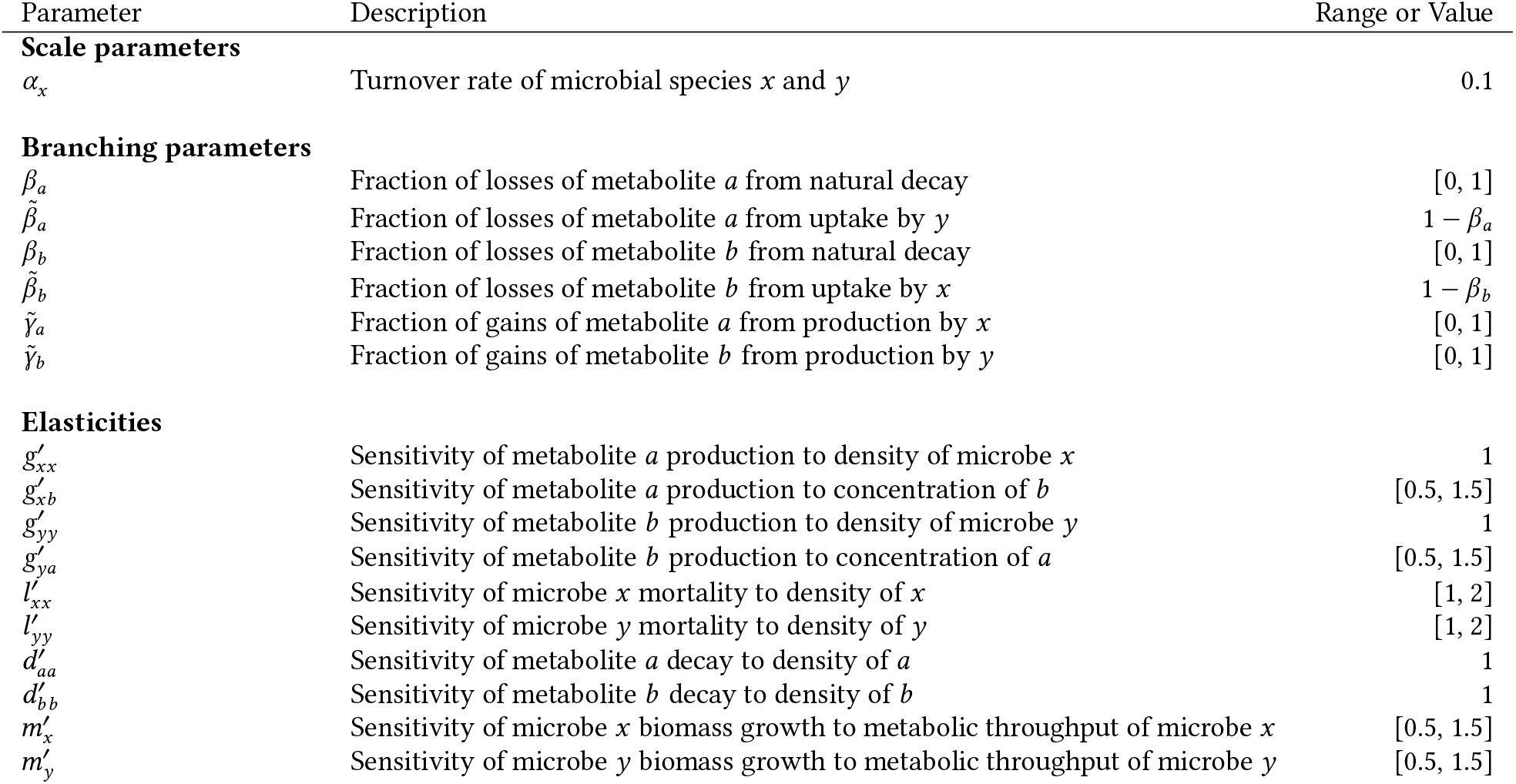
Parameter descriptions and ranges used for numerical analyses.

| Parameter | Description | Range or Value |
| --- | --- | --- |
| **Scale parameters** |  |  |
| $\alpha_x$ | Turnover rate of microbial species $x$ and $y$ | 0.1 |
| **Branching parameters** |  |  |
| $\beta_a$ | Fraction of losses of metabolite $a$ from natural decay | $[0, 1]$ |
| $\tilde{\beta}_a$ | Fraction of losses of metabolite $a$ from uptake by $y$ | $1 - \beta_a$ |
| $\beta_b$ | Fraction of losses of metabolite $b$ from natural decay | $[0, 1]$ |
| $\tilde{\beta}_b$ | Fraction of losses of metabolite $b$ from uptake by $x$ | $1 - \beta_b$ |
| $\tilde{\gamma}_a$ | Fraction of gains of metabolite $a$ from production by $x$ | $[0, 1]$ |
| $\tilde{\gamma}_b$ | Fraction of gains of metabolite $b$ from production by $y$ | $[0, 1]$ |
| **Elasticities** |  |  |
| $g'_{xx}$ | Sensitivity of metabolite $a$ production to density of microbe $x$ | 1 |
| $g'_{xb}$ | Sensitivity of metabolite $a$ production to concentration of $b$ | $[0.5, 1.5]$ |
| $g'_{yy}$ | Sensitivity of metabolite $b$ production to density of microbe $y$ | 1 |
| $g'_{ya}$ | Sensitivity of metabolite $b$ production to concentration of $a$ | $[0.5, 1.5]$ |
| $l'_{xx}$ | Sensitivity of microbe $x$ mortality to density of $x$ | $[1, 2]$ |
| $l'_{yy}$ | Sensitivity of microbe $y$ mortality to density of $y$ | $[1, 2]$ |
| $d'_{aa}$ | Sensitivity of metabolite $a$ decay to density of $a$ | 1 |
| $d'_{bb}$ | Sensitivity of metabolite $b$ decay to density of $b$ | 1 |
| $m'_x$ | Sensitivity of microbe $x$ biomass growth to metabolic throughput of microbe $x$ | $[0.5, 1.5]$ |
| $m'_y$ | Sensitivity of microbe $y$ biomass growth to metabolic throughput of microbe $y$ | $[0.5, 1.5]$ |

Using this parameterization (see *Methods* for details), the Jacobian matrix becomes:

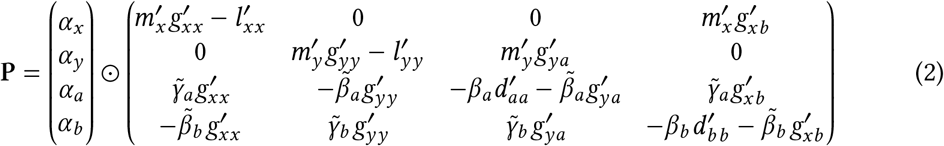

where ⊙ denotes the row-wise product, meaning that each row *i* of the matrix is multiplied by the corresponding scalar *α*_*i*_. All other parameters are defined in Table 1. A steady state is stable if all eigenvalues of the Jacobian have negative real parts.

#### What promotes stability of bacterial cooperation?

To capture the broad diversity of bacterial cross-feeding relationships, we adopt an ensemble approach [45], exploring model behavior across a wide range of biologically plausible parameter combinations (Table 1). We generate 10^7^ random parameter sets, each corresponding to a different realization of the generalized model, and evaluate the stability of each system by computing its leading eigenvalue.

Let 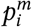 denote the *i*^th^ parameter in the *m*^th^ realization (with *i* ∈ [1, 17] and *m* ∈ [1, 10^7^]), and let *s*_*m*_ be a binary indicator of stability for parameter set *m*, defined as

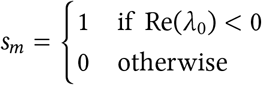

where *λ*_0_ is the dominant eigenvalue of the Jacobian. A system is considered locally stable if *s*_*m*_ = 1, and unstable otherwise. To identify which parameters promote or inhibit stability, we fit a logistic regression model (*i.e*., a generalized linear model with binomial error and logit link), using *s*_*m*_ as the binary response. All predictors are standardized to zero mean and unit variance, so the resulting coefficients reflect the direction and relative strength of each parameter’s effect on the probability of stability.

We find that system stability is positively correlated with the fraction of metabolite uptake by bacteria 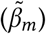, and negatively correlated with the fraction of metabolite production by bacteria 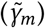 (Fig. 2A). In particular, increasing bacterial uptake of metabolite 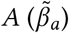 and reducing bacterial production of 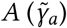 yields the highest proportion of stable systems (Fig. 2B). Overall, the ratio of uptake to production plays a key role in determining system stability.

**Figure 2:**
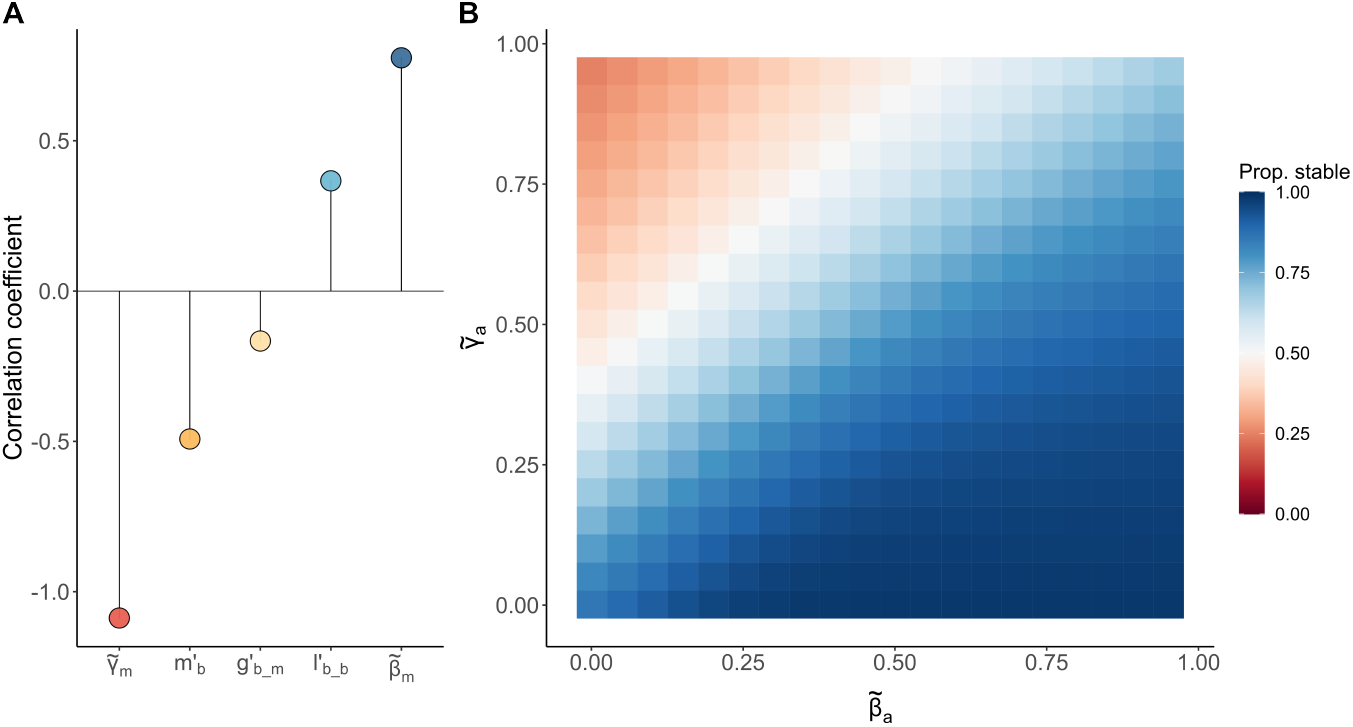
A: Correlations between model parameters and community stability, estimated as co-efficients from a binomial generalized linear model (GLM) fit to 10^7^ parameter sets (*m* = metabolite, *b* = bacteria). The shown parameters correspond to the submodel with the lowest AIC score from a global GLM including all parameters. B: Proportion of stable systems as 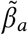 (fraction of metabolite uptake by bacterium *A*) and 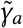 (fraction of metabolite production by bacterium *A*) are varied, based on 4 × 10^7^ parameter sets. All other parameters were drawn from uniform distributions (Table 1).

We also observe that higher sensitivity of bacterial mortality to population size 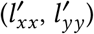 is associated with greater stability, consistent with expectations: nonlinear mortality terms (*e.g*., quadratic loss) tend to dampen fluctuations [46]. In contrast, systems become less stable when bacterial growth is highly sensitive to metabolic throughput (higher 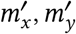), or when metabolite production is highly sensitive to the concentration of the consumed trade metabolite (higher 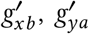). These patterns suggest strong positive feedback loops: if bacterial growth sharply increases with metabolic flux, then greater production of a metabolite promotes partner growth, which in turn accelerates production. Similarly, if metabolite production responds steeply to the concentration of the consumed metabolite, mutual amplification between partners can emerge, destabilizing the system.

#### Costly metabolite production is stabilizing

To explore patterns of stability in the bacteria-metabolite system in detail, we search for bifurcations: points in parameter space where the system undergoes qualitative changes in dynamics. Saddle-node bifurcations are particularly informative, as they mark the loss or emergence of steady states through the collision of stable and unstable branches. These bifurcations often underlie tipping points, runaway growth, or collapses in mutualistic systems [47, 48]. Identifying their locations in parameter space allows us to understand the boundaries of stability and the mechanisms by which mutualism can destabilize or persist.

Mathematically, a saddle-node bifurcation occurs when the Jacobian of the system has a zero eigenvalue. Because the determinant of a matrix equals the product of its eigenvalues, we can use a vanishing determinant of the Jacobian, |**P**| = 0, as a test function to detect such bifurcations [49]. By solving this condition symbolically for different parameters, we generate bifurcation surfaces that delineate the transitions between stable and unstable steady states [50].

Each point on these surfaces represents a parameter set at which the system changes stability (Fig. 3). Regions on one side of a bifurcation surface correspond to stable steady states, while the other side corresponds to instability. These bifurcation diagrams help us interpret how specific biological traits affect system dynamics.

**Figure 3:**
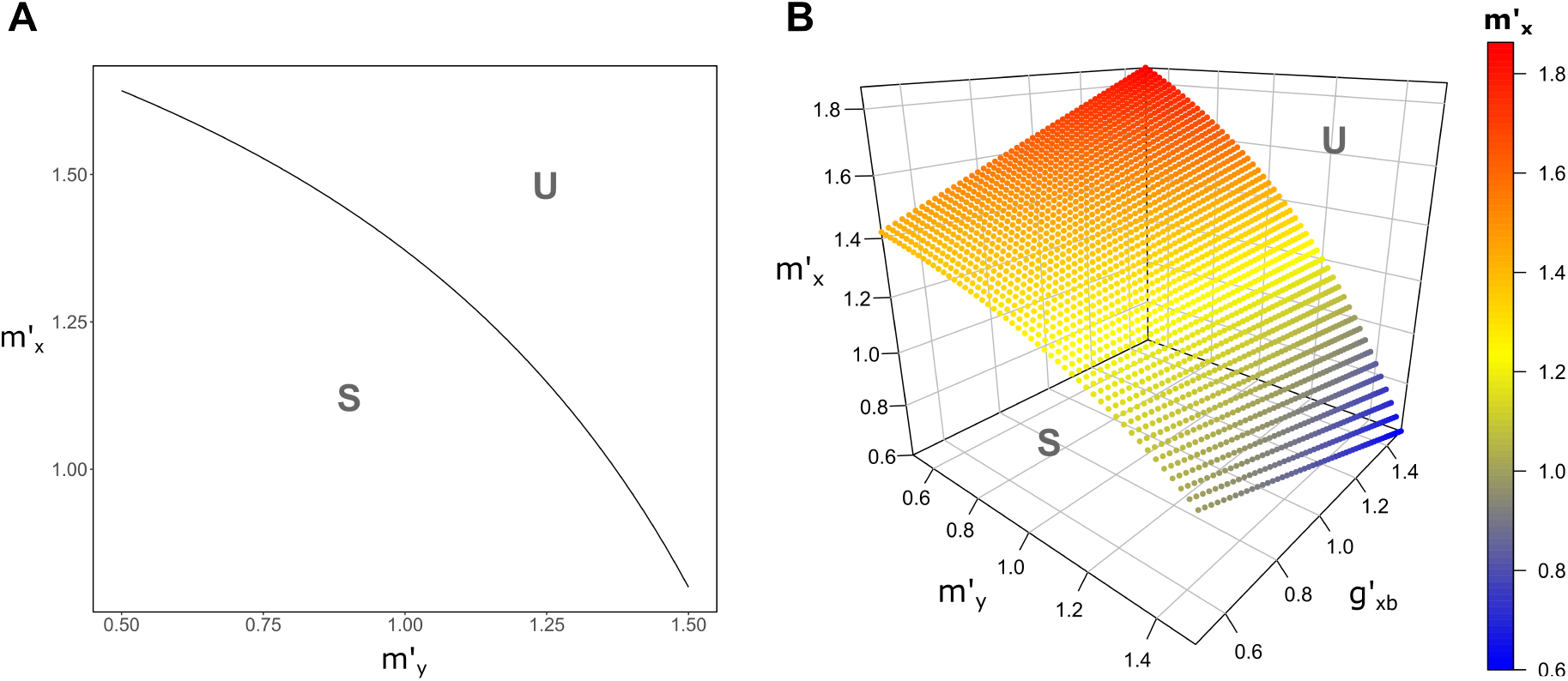
Bifurcation diagrams of bacteria-metabolite system, depending on the sensitivity of bacteria *Y* biomass growth to metabolic throughput of bacteria 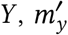, and the sensitivity of bacteria *X* biomass growth to metabolic throughput of bacteria 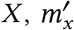, (A) and additionally on the sensitivity of metabolite *A* production to concentration of *B, g*_*x,b*_ (B). Steady states are stable (S) in the lower part/volume of parameter space. Stability is lost (U) in a bifurcation of a saddle-node type, when crossing the line/surface (parameters: 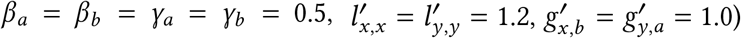.

We begin by examining the parameters 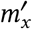 and 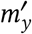, which describe how sensitively the biomass of each bacterial species depends on its metabolic throughput. Low values (< 1) reflect costly production, where bacteria grow sublinearly with increasing metabolism, for instance due to energy diverted into producing a trade metabolite like methionine [51]. High values (> 1) reflect beneficial production, where higher throughput disproportionately accelerates growth, such as when decomposing organic matter into energy.

Earlier stability analyses showed that high 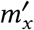 and 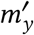 are associated with instability. The bifurcation diagram (Fig. 3A) confirms this; the system is stable in the lower-left region (low 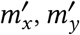) and becomes unstable via a saddle-node bifurcation as either parameter increases. Stability can be preserved if increased production efficiency in one species is counterbalanced by greater costliness in the other. This tradeoff illustrates that costly metabolite production plays a stabilizing role in cross-feeding mutualisms by dampening runaway feedbacks.

In contrast, when both partners produce metabolites cheaply, each benefits more from the other’s abundance, reinforcing a positive feedback loop that can lead to instability. Costly synthesis weakens this loop, making mutualistic growth self-limiting rather than self-reinforcing.

To understand how perturbations drive the system away from equilibrium at these bifurcations, we examine the eigenvectors associated with the eigenvalue that crosses zero. These vectors indicate the initial direction of destabilization. At the bifurcation, both bacterial species and one metabolite increase or decrease together, while the other metabolite shifts oppositely. Notably, the population of the species with cheaper metabolite synthesis exhibits larger changes, indicating that it dominates the dynamics near the tipping point.

We next examine the parameter 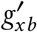, which determines how the concentration of metabolite *B* affects the production of trade metabolite *A* by species *X*. Low values (< 1) represent saturation effects — *e.g*., when production maxes out despite increasing *B* due to trace requirements or limited uptake machinery. High values (> 1) indicate enhanced production with increasing *B*, as in enzyme induction responses to lactose in *E. coli* [52].

In Fig. 3B, we plot bifurcations in the space of 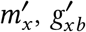, and 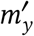. The effect of 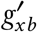 depends on the costliness of metabolite synthesis by species *Y*: when *Y* ‘s synthesis is costly, higher 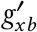 is stabilizing, while for cheap synthesis, lower 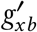 becomes stabilizing. Thus, the sensitivity of metabolite production to partner abundance can either reinforce or buffer feedback loops, depending on whether trade is energetically costly.

However, the stabilizing or destabilizing effect of 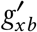 is smaller than that of 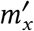 or 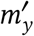. As before, eigenvector analysis shows that at the bifurcation, both bacterial populations and one metabolite change in tandem, with the species producing metabolites more cheaply driving the shift.

#### Ratio of consumption to production as a key factor in stability

We now shift our focus to the role of branching parameters in determining system stability. The fractions of metabolite supply to the system, *γ*_*a*_ and *γ*_*b*_, together with the sensitivity of bacterial *X* biomass growth to its metabolic throughput, *m* ^′^ *x*, define a three-dimensional parameter space (Fig. 4). We plot bifurcation diagrams for three distinct values of the fractions of natural metabolite decay, *β*_*a*_ and *β*_*b*_: low fractions (0.1) in panel (A), intermediate fractions (0.5) in panel (B), and high fractions (0.9) in panel (C). Generally, increasing the external supply of metabolites to the system (*γ*_*a*_ and *γ*_*b*_) stabilizes the system, while increasing the decay fractions (*β*_*a*_ and *β*_*b*_) destabilizes it.

**Figure 4:**
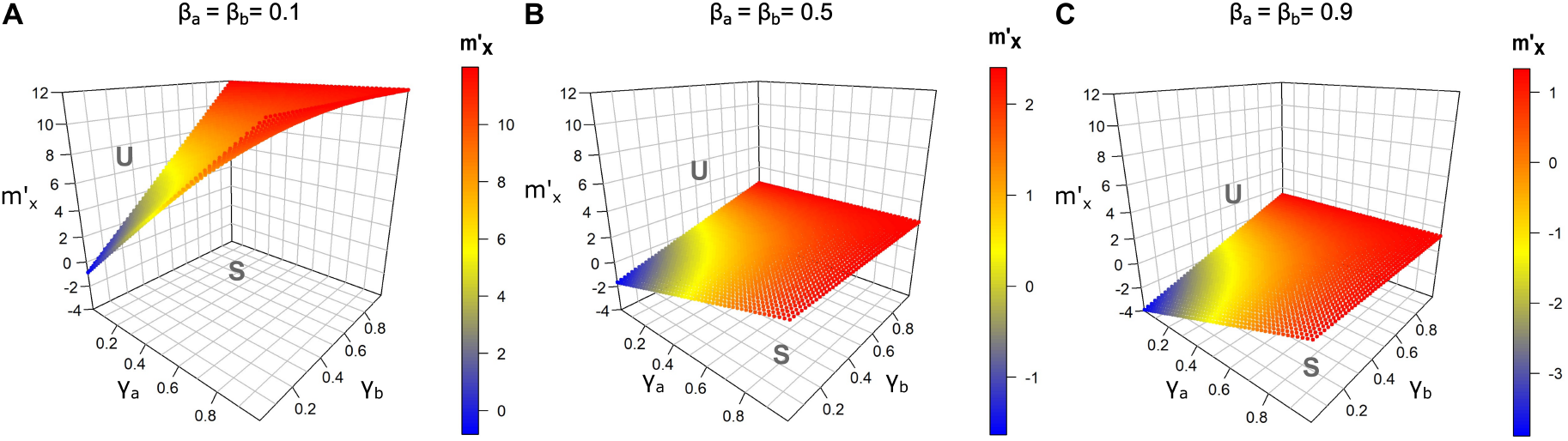
Bifurcation diagrams of bacteria-metabolite system, depending on the fraction of supply of metabolite *A, γ*_*a*_, fraction of supply of metabolite *B, γ*_*b*_, and sensitivity of bacteria *X* biomass growth to metabolic throughput of bacteria 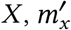. Fraction of losses of metabolite *A, β*_*a*_, and metabolite *B, β*_*b*_ are set to 0.1 (A), 0.5 (B) and 0.9 (C). Note the different color scales. Steady states are stable in the lower volume of parameter space. Stability is lost in a bifurcation of a saddle-node type, when crossing the line/surface (parameters: 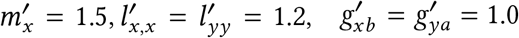.

This finding challenges the initial assumption that low bacterial metabolite production (high *γ*_*a*_ and *γ*_*b*_) and low uptake by the bacteria (high *β*_*a*_ and *β*_*b*_) would weaken the positive feed-back effect due to reduced coupling between the bacterial populations. In fact, the stability of the positive steady state hinges on the ratio of metabolites consumed by the bacteria relative to the fraction of the trade metabolite they produce. When bacteria primarily consume the partner’s metabolite and make only a minor contribution to the partner’s trade metabolite concentration, the system tends to stabilize. Eigenvector analysis of the largest eigenvalue at the bifurcation reveals that the bacterial population benefiting most from the mutualistic cross-feeding relationship experiences the greatest changes, further underscoring the stabilizing role of consumption-to-production ratios.

#### Insights from symmetric cooperation

The four-dimensional system still presents a fair amount of complexity. To gain deeper insight, we examine a special case where both species share the same set of generalized parameters and similarly for the two metabolites. This symmetry allows us to reduce the system to two simpler, two-dimensional systems. The advantage of this reduction is that it provides a more manageable framework for analysis, enabling us to extract analytical results and a clearer understanding of the system’s dynamics and stability.

To achieve this reduction we take advantage of the structure of the Jacobian of the 4-dimensional system. Assuming equal steady state values of the two bacterial populations and the two metabolite concentrations respectively as well as equal parameter pairs, e.g. 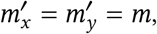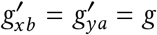, we see that in this case, the matrix

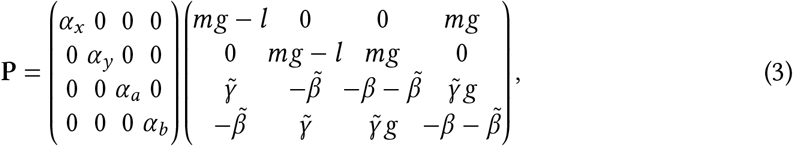

is of the structure

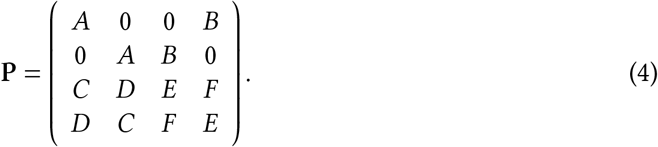

Now that the matrix has been reduced to a relatively simple structure, one can try to identify the structures of eigenvectors that are compatible with this structure. The present *P* breaks into four blocks where each block has a symmetry such that the structure of the matrix remains unchanged if the numbering of the two populations and two metabolites is reversed. Due to this symmetry all eigenvectors must have the structure (*a, a, b, b*) or (*a*, −*a, b*, −*b*), where *a* and *b* are constants that still need to be determined.

By multiplying these structures for the eigenvectors with the matrix we arrive at a condition that constrains the values of *a* and *b*. For example

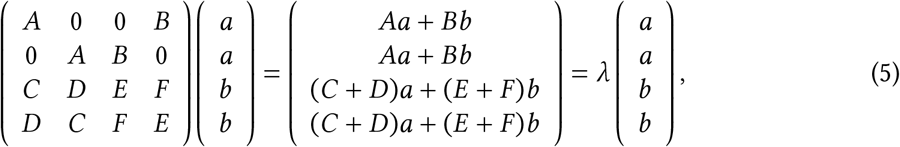

where the second equality is true if the vector is the eigenvector corresponding to *λ*. This second equality relates linear expressions of two unknowns (*a, b*) and hence can be written in the more compact two-dimensional form

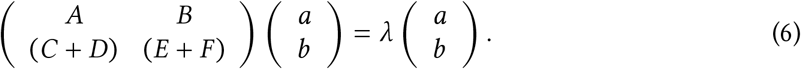

We can therefore find two eigenvalues *λ* of *P* by solving the two-dimensional eigenvector equation Eq. 6.

Similarly,

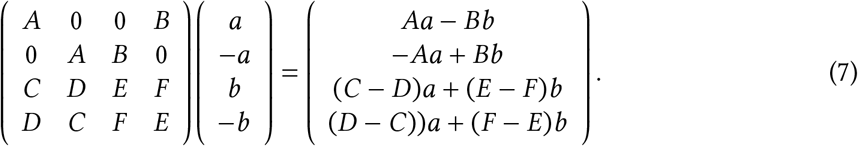

In this case we get

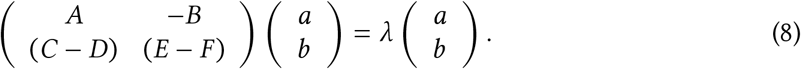

While in the first case the instabilities obey the symmetry, in this case they break the symmetry.

Taken together, we expect to get two eigenvectors from the symmetric case and two from the anti-symmetric one. This approach enables us to identify all eigenvectors of the 4 × 4 matrix. Since there are no additional eigenvectors, *λ* can only be an eigenvalue of the 4 × 4 matrix if it is an eigenvalue of at least one of the 2 × 2 matrices. Consequently, assuming equal steady state values of the bacteria and the metabolites and equal parameter pairs, we can write the 4 × 4 Jacobian of the 4-dimensional system as two 2 × 2 Jacobians

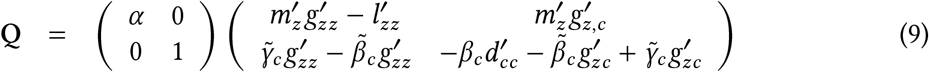

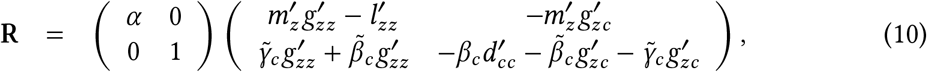

where *z* now represents the bacteria and *c* denotes the metabolites. In the following, we set 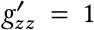 and 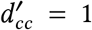 as before and use a simplified notation, i.e. 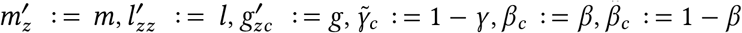.

The simplicity of the eigenvalue analysis for 2 × 2 matrices enables the derivation of stability conditions for the two-dimensional systems. In such a system, a steady state is stable if the determinant of the Jacobian is positive and the trace is negative. These conditions are indicative of the system returning to equilibrium following small perturbations. At a Hopf bifurcation, the trace becomes positive, signaling the onset of oscillatory behavior. In contrast, a saddle-node bifurcation occurs when the determinant becomes negative, reflecting a loss of stability through the merging or annihilation of steady states.

To determine whether a Hopf bifurcation can occur under symmetric conditions, we consider

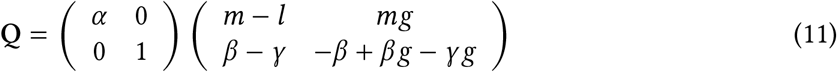

with

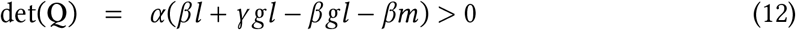

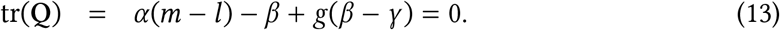

We know that −*β* < 0 and *g*(*β* − *γ*) < 0 if *β* < *γ*, hence *β* and *γ* have to take small values and *m* > *l* for a Hopf bifurcation to occur.

For the Jacobian of the anti-symmetric case we consider

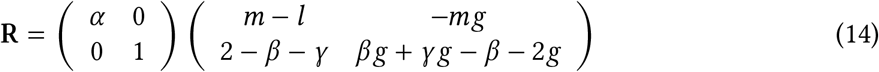

With

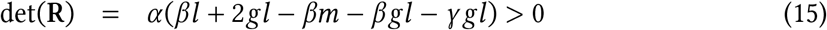

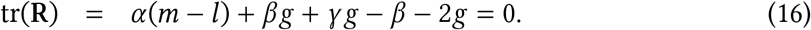

A Hopf bifurcation within a realistic parameter range will not occur in the anti-symmetric case, because the condition *βg* + *γ g* < 2*g* ensures that the trace of the Jacobian always remains negative. Hence, in a symmetric case characterized by tight cooperation between the two bacterial species, periodic cycles between the bacteria and the metabolites could occur, driven by resource exploitation. We however do not expect periodic cycles of shifting dominance between the two species.

For the saddle-node type bifurcations, we consider

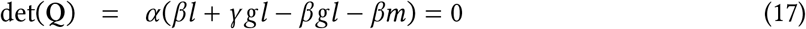

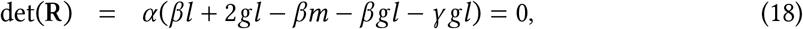

while keeping in mind that the trace values have to remain negative for the saddle-node type bifurcation to occur in the two-dimensional systems. To better understand the stabilizing or destabilizing effects of individual parameters, we take the derivative of the determinant of the Jacobian with respect to each parameter.

First, we examine the Jacobian **Q**, corresponding to the scenario where the instabilities satisfy the symmetric condition. We derive

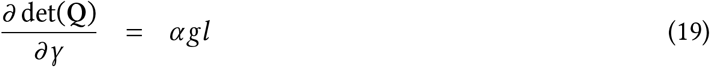

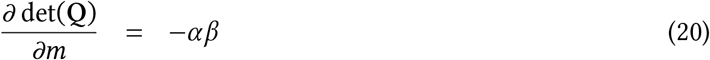

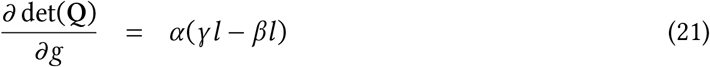

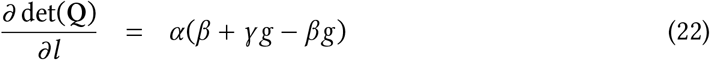

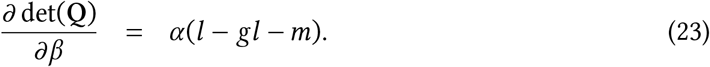

As expected, we see that *γ* consistently has a stabilizing effect, whereas *m* is always destabilizing. Additionally, we see again that the impact of *g* is influenced by the ratio of *γ* to *β*. Specifically, when *γ* > *β*, higher values of *g* contribute to system stabilization. Conversely, when *γ* < *β*, higher values of *g* lead to destabilization.

Interestingly, the term *g*(*β* − *γ*) − *β* which appears in the derivative with respect to *l*, is a component of the trace. For *l* to be destabilizing, we require *g*(*β* −*γ*) −*β* > 0, which implies that *β* > *γ* and *g* > 1. For the trace to remain negative, we also need *l* > *m*. Therefore, increasing *l* can destabilize a system, where the bacteria are highly dependent on the production of the partner’s metabolite, but derive limited benefits from producing the trade metabolite. However, such systems are not stable within the plausible parameter range. Additionally, increasing the parameter *β* is destabilizing if *m* > 1 or *g* ≥ 1.

Next, we consider the Jacobian **R**, corresponding to the scenario where the instabilities violate the symmetric condition. We derive

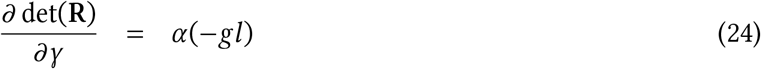

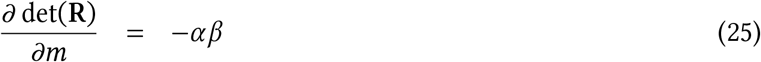

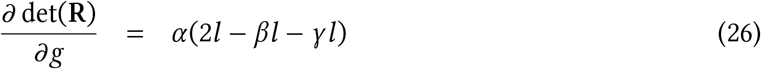

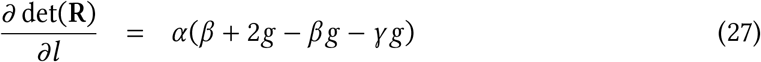

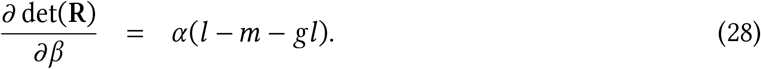

The conditions for *m* and *β* in this scenario mirror those found for the Jacobian **Q**. In the considered parameter range, large values of *g* and *l* consistently have a stabilizing effect, since 2 > *γ* + *β*. In this case, *γ* acts as a destabilizing factor. This may be because an increase in *γ* reduces the bacteria’s dependence on the interaction, potentially leading to divergent trajectories for the two bacterial populations.

Intriguingly, the effect of *γ*, the fraction of metabolite supply to the system, strongly differs between the different scenarios, even in this simple model. This could have important implications for the use of prebiotics and, more broadly, in medicine, as treatment outcomes may depend strongly on a multitude of other parameters, such as the tightness of cooperation or the coupling of one metabolite’s concentration to the other metabolite’s production.

Investigating the impact of a small additional loss of metabolites on the system’s steady state,

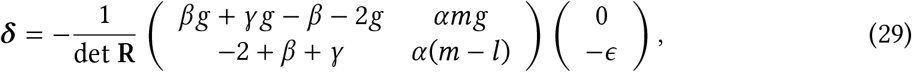

results in

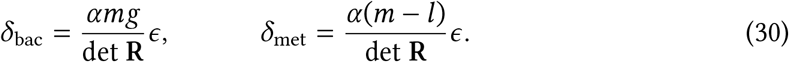

Interestingly, a loss of metabolites leads to an increase in bacterial density. This may happen in the anti-symmetric case because decreasing the metabolite concentration increases the dependence on the trade partner’s metabolite production, making a shift in dominance of one bacterium less likely. The response of the metabolite concentration depends on the ratio *m*/*l*. If *m* > *l*, then the loss in metabolite results in an increase in metabolite concentration and vice versa.

In this section, we have seen how different the effects of the parameters and changes in the system can be, depending on which instability we consider. While in the symmetric case, the independence from the trading partner’s production stabilizes the system, in the anti-symmetric case, the opposite is true. This can lead to interesting observations, such as an increase in bacterial populations in response to a decrease in metabolites — a counter-intuitive response, also known as a hydra effect [53].

### Impacts of perturbations to bacteria and metabolites

We next explore the impact of press perturbations (*i.e*., permanent changes to parameters) on the system using a linear approximation. To quantify the net impact of perturbations among bacteria and metabolites, we account for both direct and indirect effects through all paths in the motif [54, 55, 56, 57, 58], which can be directly derived from the Jacobian. For example, we consider how the system responds to a fixed decrease in the bacterial population *X* or a drop in metabolite *A* concentration as other bacteria begin to consume the metabolite. These are linearized representations, not exact reductions.

Conceptually, we compute the steady state of the system as a function of parameters *s*, written as **x**^⋆^ = *x*^⋆^(*s*). Since 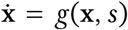 and *g*(**x**^⋆^(*s*), *s*) = 0, we differentiate with respect to *s* using a corollary of the implicit function theorem [59, 57], yielding:

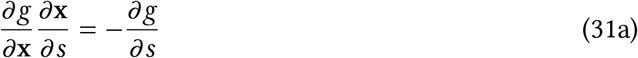

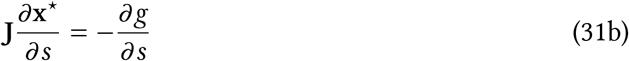

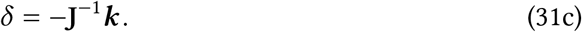

where ***δ*** is the steady-state shift, **J**^−1^ is the Jacobian matrix inverse, and ***K*** = ∂*g*/∂*s* is the vector describing the direct impacts of the perturbation on the equations [57]. For example, the direct effect of decreasing the bacteria *X* population by a small quantity might be represented as

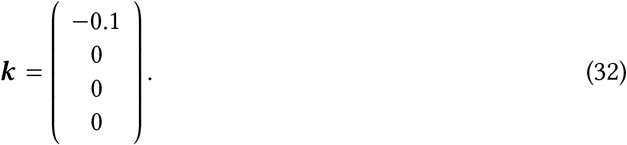

The impact vector ***δ*** captures the indirect relative changes in bacteria or metabolite levels after the perturbation propagates through the motif. These changes are normalized relative to the unperturbed steady state and expressed per unit of direct effect. Negative values indicate losses, positive values represent gains, and larger absolute values indicate more significant impacts.

We calculate ***δ*** for ∼ 7 × 10^6^ parameter sets, excluding unstable states, and apply Eq. 31c for different perturbation scenarios (Fig. 5). Across all scenarios, a consistent pattern emerges: the impact is greatest for at least one bacterial species. Notably, altering bacterial density has a much stronger effect (up to 7.5 times greater) on the system than changing metabolite concentrations. Shifts in metabolite levels primarily influence the bacterial species that consumes those metabolites, while changes in bacterial population density affect the corresponding bacterial population. These changes also exert a smaller, opposing effect on the metabolites consumed by the affected bacterial species. Overall, our results demonstrate that variations in bacterial density have greater leverage on the mutualistic cross-feeding system than changes in metabolite concentrations.

**Figure 5:**
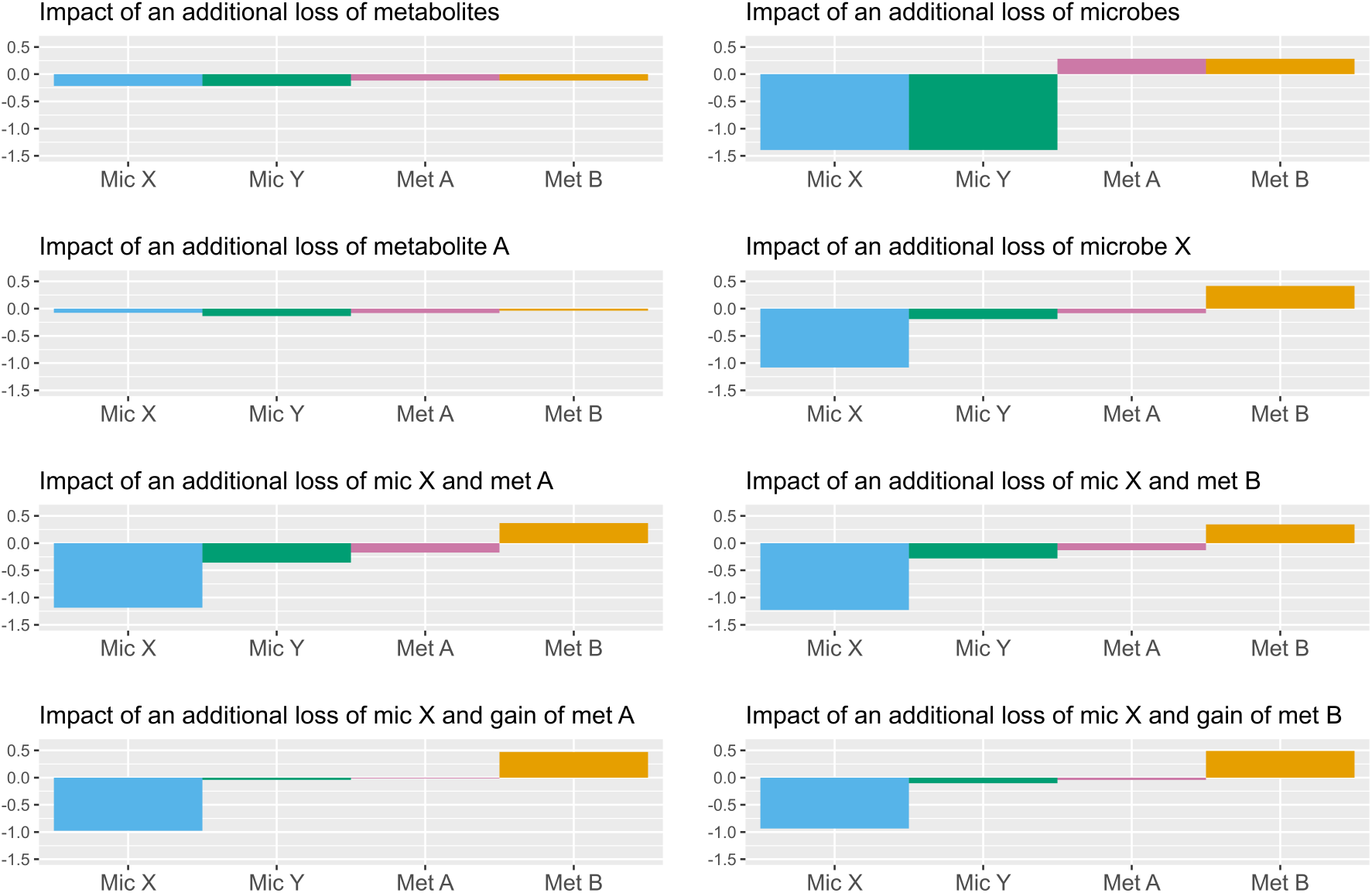
Impact of press perturbations (*e.g*., a small decrease in bacterial or metabolite density) on the cross-feeding motif. The y-axis shows the negative or positive impact on the system. Each bar represents the median impact for each bacterium (mic) and metabolite (met) type, calculated from ∼ 7 × 10^6^ parameter sets.

### Dynamics of spatial trade networks

The cross-feeding motif we’ve analyzed so far often exists within broader spatial trade networks in nature [9], where metabolite exchange is not just local but distributed across space [60]. These spatial dynamics can shift the balance of cooperation and competition, and drive pattern formation at scales much larger than an individual patch.

To explore this, we embed the motif (Fig. 1) within spatial networks and examine the emergence of diffusion-driven Turing instabilities [61]. These arise when the diffusion of metabolites interacts with local feedbacks in a way that destabilizes spatially homogeneous equilibria, producing patterned distributions across habitat patches. Such instabilities can generate persistent heterogeneity in microbial systems [62, 63].

We extend Eq. 2 to a spatial network using the master stability function approach [64]. In this framework, each node of the network contains an identical copy of the local interaction motif, and metabolites diffuse along links between nodes. The meta-Jacobian of the full system can be written as

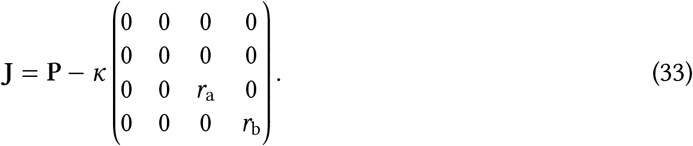

Here, *κ* is an eigenvalue of the Laplacian matrix describing the spatial network topology, and **C** (diagonal in this case) contains the metabolite diffusion rates *r*_a_ and *r*_b_. This formulation assumes that metabolites diffuse on much faster timescales than microbial growth or movement. When *κ* = 0, the system reduces to the non-spatial case. The key insight is that spatial structure can only destabilize a system that is already stable in the absence of diffusion and never the reverse. This follows because the Laplacian is positive semidefinite, so it always has a zero eigenvalue corresponding to the homogeneous case. If any nonzero *κ* pushes an eigenvalue of **J** into the right half-plane, the system becomes unstable to spatial perturbations.

Figure 6 illustrates an example. Panel A shows the master stability function, which relates the system’s Jacobian eigenvalues to the Laplacian spectrum [64, 60]. The blue region marks the instability band; if any Laplacian eigenvalue falls in this interval, spatial patterns emerge. Increasing the global coupling strength (bubble size in Fig. 6A) stretches the Laplacian spectrum and increases the risk of instability.

**Figure 6:**
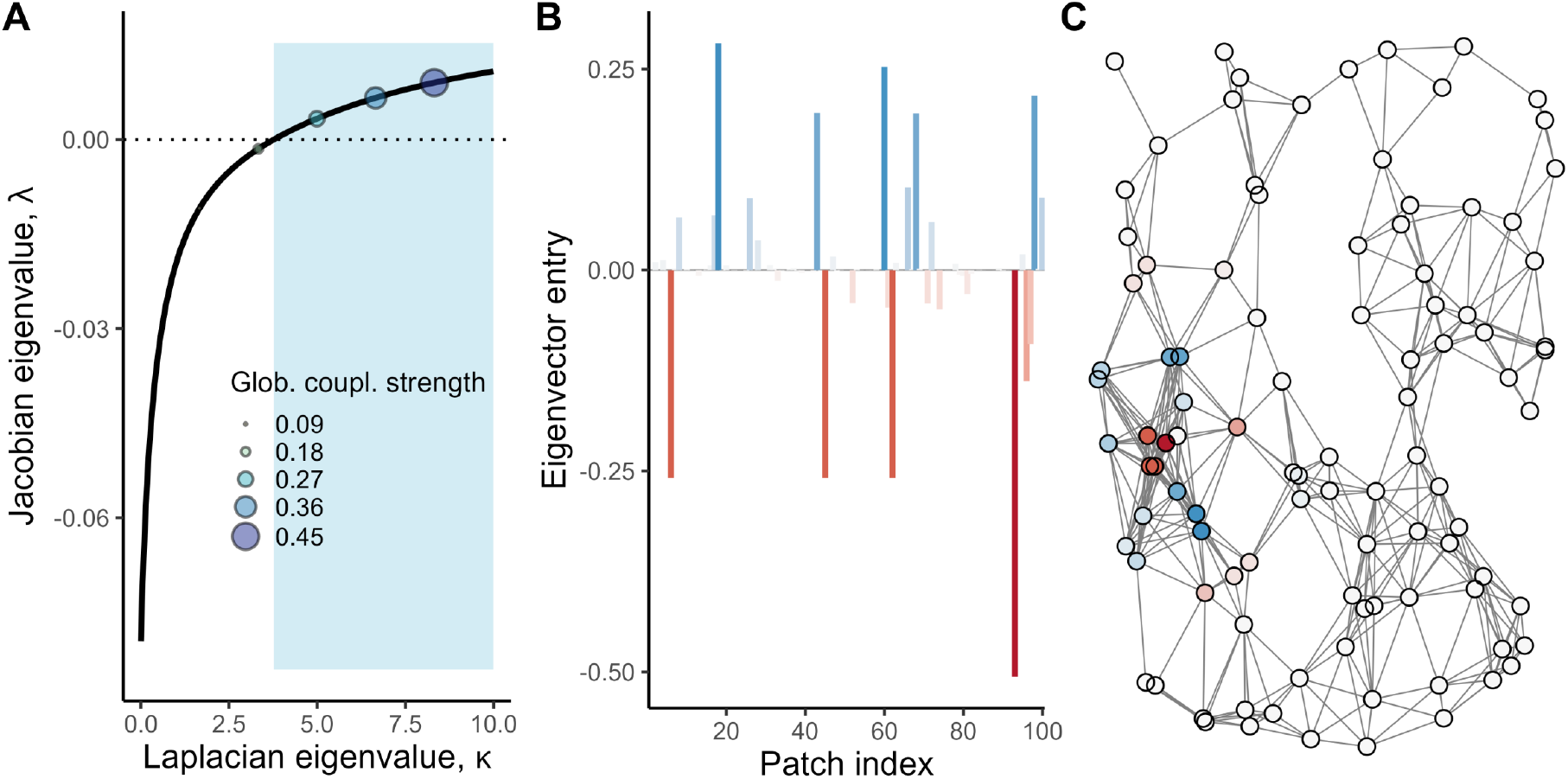
Example of a diffusion-driven Turing instability that occurs when metabolites diffuse. The master stability function (MSF) (A) links the Jacobian eigenvalue of the spatial system to the Laplacian eigenvalue of the spatial network. The spatial system is stable if none of the Laplacian eigenvalues lie within the interval where the MSF takes on positive values (blue shaded area). Increasing global coupling strength (Glob. coupl. strength) can destabilize the system by expanding the range of the Laplacian spectrum. Panel B shows the eigenvector entries of the different patches and panel C shows the example of a fully connected random geometric graph with a 100 nodes and a radius of 0.32, where the patches are colored according to their eigenvector entries corresponding to microbe *X*.

Interestingly, the resulting instability is highly localized (Fig. 6B and C), affecting only a few nodes despite occurring in a large, well-connected network. This localization arises from the structure of the Laplacian eigenvectors and the nonlinear feedback loop in the system, where bacterial growth amplifies local increases in metabolite concentration, leading to spatially heterogeneous patterns and spatial reciprocity.

## Discussion

Using a generalized modeling approach, in this work we explored a simple cross-feeding motif commonly observed in bacterial systems. By focusing on the general structure of the system rather than specific functional forms, our model enabled an extensive exploration of cross-feeding dynamics. This included transitions from the release of inexpensive metabolites to the synthesis of expensive ones, as well as scenarios where bacteria were either the predominant contributors to metabolites or contributed only marginally to the overall metabolite pool. The low-dimensionality of our model allowed for a comprehensive examination of key qualitative features underlying cross-feeding dynamics.

Our findings underscore the critical role of metabolic costs associated with metabolite production in determining system stability. The exchange of inexpensive metabolites tends to introduce instability, while the production of costly metabolites emerges as a stabilizing factor. Notably, systems involving the low-cost production of one metabolite can achieve stability when paired with high-cost production of a complementary metabolite in the cross-feeding relationship. An illustrative real-world example is the interaction between acetate-releasing *E. coli*, which produces metabolites with low metabolic costs, and methionine-releasing *S. enterica*, where production is metabolically expensive [51].

Patterns across sequenced bacterial genomes also support this framework. Cooperative interactions involving high-cost amino acids have been found to be more robust than those involving cheaper compounds [10]. An analysis of over 6,000 bacterial genomes showed that only a small subset of taxa produce the most metabolically expensive amino acids, while the majority rely on these key producers. This uneven distribution suggests that communities may be structured around the limiting availability of costly metabolites, in much the same way that environmental nutrients regulate growth in classical ecological systems [65].

Another key determinant of stability in our model is the ratio between the fraction of trade metabolites taken up by bacteria and the fraction they produce. When uptake exceeds production *i.e*., when this ratio is greater than one, stability is enhanced, likely due to the weakening of the system’s inherent positive feedback. This happens because bacteria are absorbing more metabolite for their own use than they are contributing to their partner’s metabolite pool. In real-world microbial systems, this balance can shift depending on the external nutrient supply, which in turn can transform the nature of the interaction from mutualism to competition, or even to competitive exclusion [47]. At low nutrient concentrations, cooperation becomes more favorable, as partner dependency increases [47]. These dynamics reflect the broader reality that microbial interactions are not fixed but highly context-dependent, varying across environmental conditions, spatial structure, and time [66].

When examining the nature of instabilities, a recurring pattern emerged in which one species consistently showed stronger responses than its partner. This asymmetry suggests that instabilities often involve a single species dominating system behavior, potentially due to becoming more abundant. These shifts are shaped by differences in nutrient supply rates, uptake efficiencies, and metabolic costs, leading to directional changes in abundance rather than synchronized rises or declines. The non-trivial nature of these instabilities highlights how nuanced variations in parameter values can produce complex dynamic outcomes.

Earlier models of mutualism, particularly those abstracting away resource dynamics, tended to view these interactions as inherently destabilizing due to their positive feedback structure [19, 20]. However, more recent models that explicitly incorporate resource exchange have shown that mutualism tends to remain unstable unless interaction strengths are weak or partners reciprocate with high fidelity [7, 8]. In contrast, the framework developed here shows that the metabolic costs of producing shared metabolites can act as a counterbalancing stabilizing force. As discussed above, stability is further enhanced when costly metabolites are exchanged, potentially offering an evolutionary pathway toward robust cooperative interactions in microbial systems [67].

Beyond identifying factors that shape the stability of steady states in cross-feeding systems, we also examined how the system responds to perturbations. Overall, changes in bacterial density had a more pronounced effect on system behavior than changes in metabolite concentrations. Across all perturbation types, the bacterial components proved more influential than the metabolites themselves. This asymmetry has practical implications, particularly in efforts to manipulate microbial communities, for instance, using pro-or prebiotics to shift bacterial composition or function [68]. From an applied perspective, our model suggests that probiotics may be more effective, in general, at modifying the structure of bacterial communities with cooperative interactions.

Extending the analysis to spatially structured systems, we explored a network of homogeneous patches and found that while the basic cross-feeding motif remains stable in isolation, the introduction of metabolite diffusion can destabilize the homogeneous state and lead to spatial pattern formation. Stronger diffusive coupling widens the Laplacian spectrum, increasing the potential for Turing-like instabilities in a networked context [64, 60]. Though our model did not produce spatial patterns in a continuous domain, the framework offers a foundation for future investigations into spatially extended microbial dynamics. The emergence of cross-feeding motifs across space allows bacteria to build distributed metabolic networks, effectively sharing metabolic functions and shifting metabolism from a cell-autonomous trait to a communal one [3]. These spatial patterns can be critical for the survival of microbial communities [69], influence their evolutionary trajectories [70], and shape emergent community-level properties [71].

## Conclusions

This generalized modeling framework for microbial cooperation reveals a rich set of dynamics, highlighting when steady states are stable, how the system responds to disturbance, which components are most influential, and how spatial processes modify outcomes. The value of the generalized modeling approach lies in its flexibility: it accommodates a wide range of functional forms and parameter spaces while remaining analytically tractable. This accessibility makes it especially useful for generating quick, interpretable insights into microbial behavior and for building intuition about the principles underlying cooperative interactions in complex biological systems.

## Methods

To parameterize the Jacobian in an interpretable way, we assume that all variables and process rates have positive values. Furthermore we observed that in the space of plausible functions many steady states must exist. In the following we now study the stability of all of these steady states. We denote these steady states values of variables as *X*^*^ and and the rates of process in the steady state as *P* ^*^ = *P* (*X*^*^) and normalize the equations with respect to *X*^*^. For every variable *X* we define

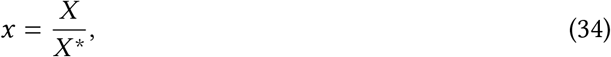

and for every process *P* (*X*) we define

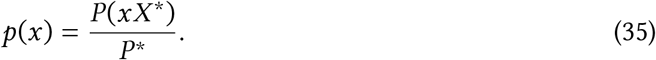

For the growth rates this means

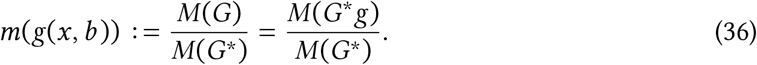

This leads to the normalized differential equations

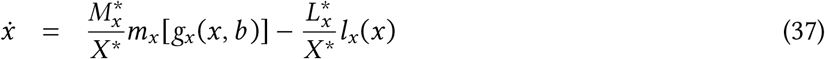

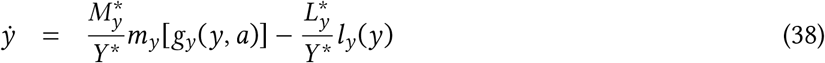

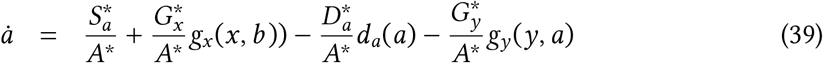

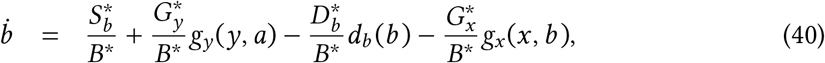

where normalized functions are indicated by lowercase letters. This normalization procedure maps the unknown state to a known location *x*^*^ = *y*^*^ = *a*^*^ = *b*^*^ = 1, and in the steady state all processes run at rate 1. Because gains and losses are balanced in the steady state, we can define scale parameters representing metabolite and microbe turnover rates

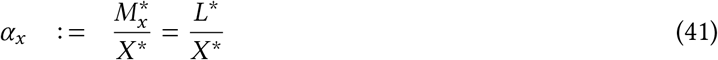

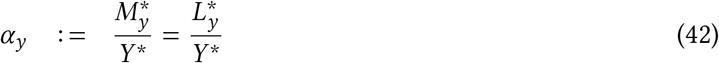

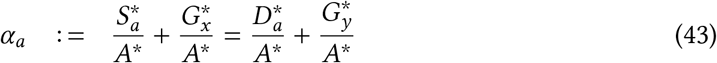

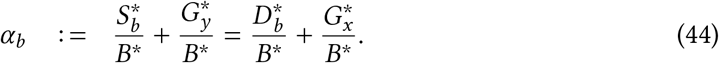

To quantify the relative contribution of each gain and loss process to population turnovers we define branching parameters [42, 43]

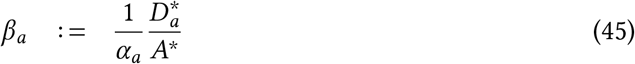

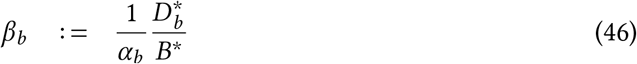

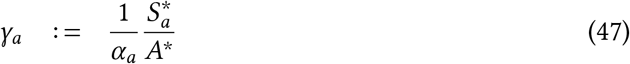

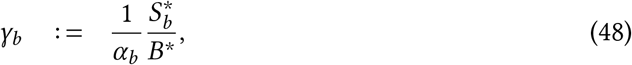

and the complementary parameters 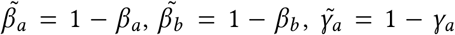 and 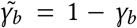. For example, when 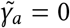, it means that all of metabolite *A* originates from an external source, indicating zero bacterial production of metabolite *A*. Conversely, if 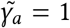, there is no external supply of metabolite *A* to the system; instead, it is entirely produced by the bacteria. When 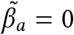, metabolite *A* undergoes decay exclusively, without any uptake by bacteria. Conversely, if metabolite *A* is solely removed from the environment through bacterial uptake, implying no natural decay, then 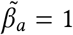.

Descriptions and ranges for scale and branching parameters are provided in Table 1 and discussed by [42, 44]. Substituting scale and branching parameters into eqs. 37-40 yields

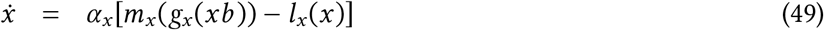

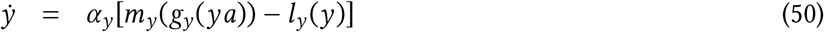

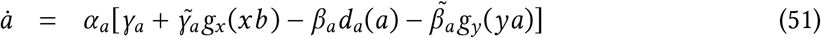

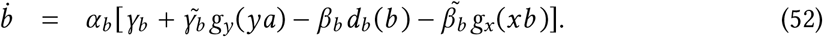

Local stability of this system can be analyzed by calculating the corresponding Jacobian matrix **P**, which determines the behavior of the system close to the steady state. It is defined by 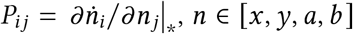. The Jacobian contains additional parameters called elasticities or exponent parameters [42, 43], where for instance

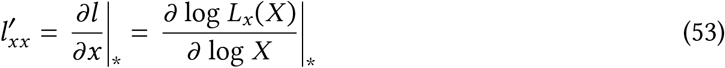

are logarithmic derivatives, and |_*_ indicates that the derivatives are evaluated at the steady state. The double subscript “*x, x*” specifies our reference to bacteria labeled as *X* and denotes that the function is undergoing differentiation with respect to *x*. Consequently, the parameter 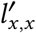 serves as an indicator of the sensitivity of bacteria *X* mortality to the density of *X*. In general these parameters are a measure of nonlinearity of the process at the steady state. Exponent parameters are defined in Table 1. We obtain the Jacobian

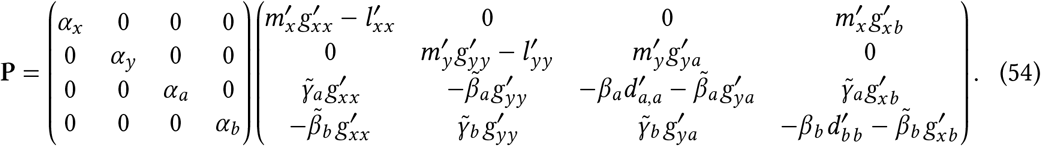

A steady state is stable if all eigenvalues of the system’s Jacobian have negative real parts, and a loss of stability occurs if the real parts of one or more eigenvalues become positive.

## Acknowledgments

We thank J.L. Green for discussions; and the Santa Fe Institute for sponsoring the “Predicting microbiome response to disturbance” working group, which inspired this work. J.C.M. was funded by the European Research Council, ERC grant, C-Quest grant no. 101044738. A.K.F was supported by Alfred P. Sloan Foundation grant no. 2017-9807 and National Science Foundation grant no. EF-2222478.

